# Specific activation of HIV-1 from monocytic reservoir cells by bromodomain inhibitor in humanized mice in vivo

**DOI:** 10.1101/375535

**Authors:** Guangming Li, Zheng Zhang, Natalia Reszka-Blanco, Feng Li, Liqun Chi, Jianping Ma, Jerry Jeffrey, Liang Cheng, Lishan Su

## Abstract

The combination antiretroviral therapy (cART) effectively suppresses HIV-1 infection and enables HIV-infected individuals to live long productive lives. However, the persistence of HIV-1 reservoir cells with latent or low-replicating HIV-1 in patients under cART make HIV-1 infection an incurable disease. Recent studies have focused on the development of strategies such as epigenetic modulators to activate and purge these reservoirs. Bromodomain inhibitors (BETi) are epigenetic modulating compounds able to activate viral transcription in HIV-1 latency cell lines in a positive transcription elongation factor b (P-TEFb)-dependent manner. Little is known about the efficacy of activating HIV-1 reservoir cells under cART by BETi in vivo. In this study, we seek to test the potential of a BETi (I-BET151) in activating HIV-1 reservoir cells under effective cART in humanized mice in vivo. We discover that I-BET151 efficiently activates HIV-1 transcription in monocytic cells, but not in CD4^+^ T cells, during suppressive cART in vivo. We further reveal that HIV-1 proviruses in monocytic cells are more sensitive to I-BET151 treatment than in T cells in vitro. Finally, we demonstrate that I-BET151-activated viral transcription in monocytic cells is dependent on both CDK2 and CDK9, whereas only CDK9 is involved in activation of HIV-1 by I-BET151 in T cells. Our findings indicate a role of myeloid cells in HIV-1 persistence, and highlights the limitation of measuring or targeting T cell reservoirs alone in terms of HIV-1 cure, as well as provides a potential strategy to reactivate monocytic reservoirs during cART.

**IMPORTANCE:** It has been reported the low level of active P-TEFb critically contributes to the maintenance of HIV-1 latency or low-replication in HIV-1 reservoir cells under cART. Bromodomain inhibitors are used to activate HIV-1 replication in vitro but their effect on activation of the HIV-1 resevoirs with cART in vivo is not clear. We found that BETi (I-BET151) treatment reactivated HIV-1 gene expression in humanized mice during suppressive cART. Interestingly, I-BET151 preferentially reactivated HIV-1 gene expression in monocytic cells, but not in CD4 T cells. Furthermore, I-BET151 significantly increased HIV-1 transcription in monocytic cells, but not in latently infected CD4 T cells, via CDK2-dependent mechanisms. Our findings suggest that BETi can preferentially activate monocytic HIV-1 reservoir cells, and a combination of latency reversal agents targeting different cell types and pathways is needed to achieve reactivation of different HIV-1 reservoir cells during cART.

## Introduction

The global implementation of cART has transformed HIV infection from a fatal disease into a manageable chronic illness, and successfully blunted HIV pandemic (1-3). HIV-1 replication can be suppressed and maintained at a level below the detection limits by current cART regimens. However, cART is still unable to cure HIV-1 infection. Drug resistance, serious non-AIDS events as well as financial and societal issues associated with life-long cART highlight the urgent need in finding a cure for the disease (4-7). Therefore, recent efforts have focused on interventions that can yield a drug-free remission of HIV-1 replication or even eradication of replication-competent HIV-1 proviruses in patients.

The obstacle to eradicate HIV-1 is the persistence of latent or low replicating HIV-1 reservoirs. Following discontinuation of cART, HIV-1 reservoirs are able to produce infectious viral particles, result in viral relapse(8-11). The molecular mechanisms associated with the maintenance and reactivation of HIV latency have been extensively investigated, including transcriptional interference, deleterious mutations in the viral genome, low levels of transcriptional activators, inadequate Tat activity, epigenetics, transcriptional repressors engagement, nucleosome positioning, mRNA splicing or nuclear export blocks as well as cellular microRNA(12, 13). It is most likely that multiple mechanisms are involved in HIV-1 reservoirs persistence. And the relative importance of these mechanisms in different cell types is remain to be determined.

The best characterized HIV-1 cellular reservoir is the long-lived resting CD4^+^ T lymphocytes harboring quiescent proviral DNA that is replication-competent upon reactivation. However, reservoirs in myeloid cells could be a problem for curative strategies targeting to resting CD4^+^ T cells only. Macrophages are susceptible to HIV-1 infection in human and animal models(14-17), and HIV-1 could be recovered from the circulating monocytes pool of patients treated with cART (18, 19). In addition, HIV-1 is able to infect and replicate in brain astrocytes and microglia cells in a restricted manner that could persist despite cART (20, 21). And recent reports demonstrated that humanized mice with myeloid cells only allow HIV-1 persistent infection in macrophages during cART in vivo(17), and integrated HIV-1 DNA can be detected in the bone marrow and spleen macrophages in humanized mice with suppressive cART(22). These findings highlight that, other than resting CD4^+^ T cells, monocytes or macrophages are of great clinical importance in terms of HIV-1 cure.

It is known that bromodomain-containing protein 4 (BRD4) competes P-TEFb and disrupts the interaction between Tat and P-TEFb, and forfeits the ability of Tat to trans-activate HIV-1 transcription(23-25). Given the important role of P-TEFb in regulating HIV gene expression, different BETi have been explored and tested to activate HIV-1 gene expression in latent models of primary CD4^+^ T cells, lymphocytic T cell lines and monocytic cell lines (26, 27). However, little is known about the therapeutic potential of BETi in activating viral replication in HIV-1 reservoirs during cART in vivo.

In this study, we tested how bromodomain inhibitor (I-BET151) affected viral replication in HIV-1 reservoir cells in vivo during suppressive antiretroviral therapy. Our results demonstrate that I-BET151 treatment leads to reactivation of HIV-1 gene expression preferentially in monocytic cells during cART, which highlights the therapeutic potential of bromodomain inhibitors to activate the unique monocytic HIV-1 reservoir cells in vivo.

## RESULTS

### HIV-1 persistence in both T and myeloid cells in NRG-hu HSC mice during cART

NRG-hu HSC mice (hu-mice) were infected with HIV_JRCSF_ and monitored for viral load and HIV-1 pathogenesis in peripheral blood until 14 weeks post-infection (wpi). Several infected animals were treated with cART in mouse diet from 4-to-12 wpi. We showed that plasma viral load declined rapidly after initiation of cART, concomitant with a recovery of peripheral blood CD4^+^ T cells (Fig. 1A and Fig. S1A). Viral load reached undetectable levels in treated mice around 3-4 weeks after cART (Fig. 1A). We kept cART on board four more weeks before withdraw at 12wpi. As observed in HIV-1 patients, plasma viral load rebounded rapidly after discontinuation of cART (Fig. 1A), correlated with a decrease of human CD4^+^ T cells in the blood (Fig. S1A). At 12 wpi before cART cessation, three mice in each group were terminated for determining HIV-1 replication and pathogenesis in lymphoid tissues. The results demonstrated that HIV-1 gag p24 was significantly detected in both CD4 T cells (huCD45^+^CD3^+^CD8^-^) and monocytic cells (huCD45^+^CD3^-^CD11c^+^CD14^+^) in spleens and BM in cART-naïve mice. Accordingly, cART abolished p24 expression in both cell populations (Fig. 1B-E). Meanwhile, the CD4^+^ T cell level in cART-treated mice was significantly recovered, whereas a marked deletion of CD4^+^ T cells was observed in cART naïve-treated mice (Fig. S1A-C). The increase of CD4^+^ T cell percentage correlated with a dramatic recovery of CD4^+^ T cell number in lymphoid tissues (Fig. S1D). As we reported previously, cART-resistant replication-competent HIV-1 reservoirs persist (Fig. S2A), though cell-associated HIV-1 RNA and DNA was remarkably decreased by cART in lymphoid tissues (Fig. S2B and C) (28, 29), and viremia rebounded upon cART cessation (Fig. 1A). However, the detection of p24 in monocytic cells could be due to their phagocytosis of HIV-1 infected CD4^+^ T cells(30). In order to investigate whether monocytic cells could be infected by HIV-1, and more importantly, play a role as HIV-1 reservoirs during cART, we purified human CD3^-^ CD11c^+^CD14^+^ monocytic cells and CD3^+^CD8^-^ (CD4) T cells from bone marrow cells in animals with undetectable HIV-1 p24 positive cells by flow cytometry. We first measured cell-associated HIV-1 DNA by real-time PCR. In addition, purified monocytic cells was stimulated with TNF-α for 24 hours and then co-cultured with MOLT-4 cells to detect replication-competent virus. Total bone marrow cells from the same animals were used as controls. The results revealed that monocytic cells harbored more HIV-1 DNA per million cells on average than CD4 T cells (Fig. 1F). Notably, replication-competent viruses could be recovered from monocytic cells (Fig.1G), suggesting monocytic cells as well as CD4 T cells harbored replication-competent HIV-1 during cART. Therefore, HIV-1 persistent infection and cART-resistant reservoirs observed in humanized mouse model resembles a situation observed in HIV-1 patients. Both CD4^+^ T cells and monocytic cells can serve as HIV-1 reservoirs, contributing to HIV-1 persistence during suppressive cART.

**FIG 1.**
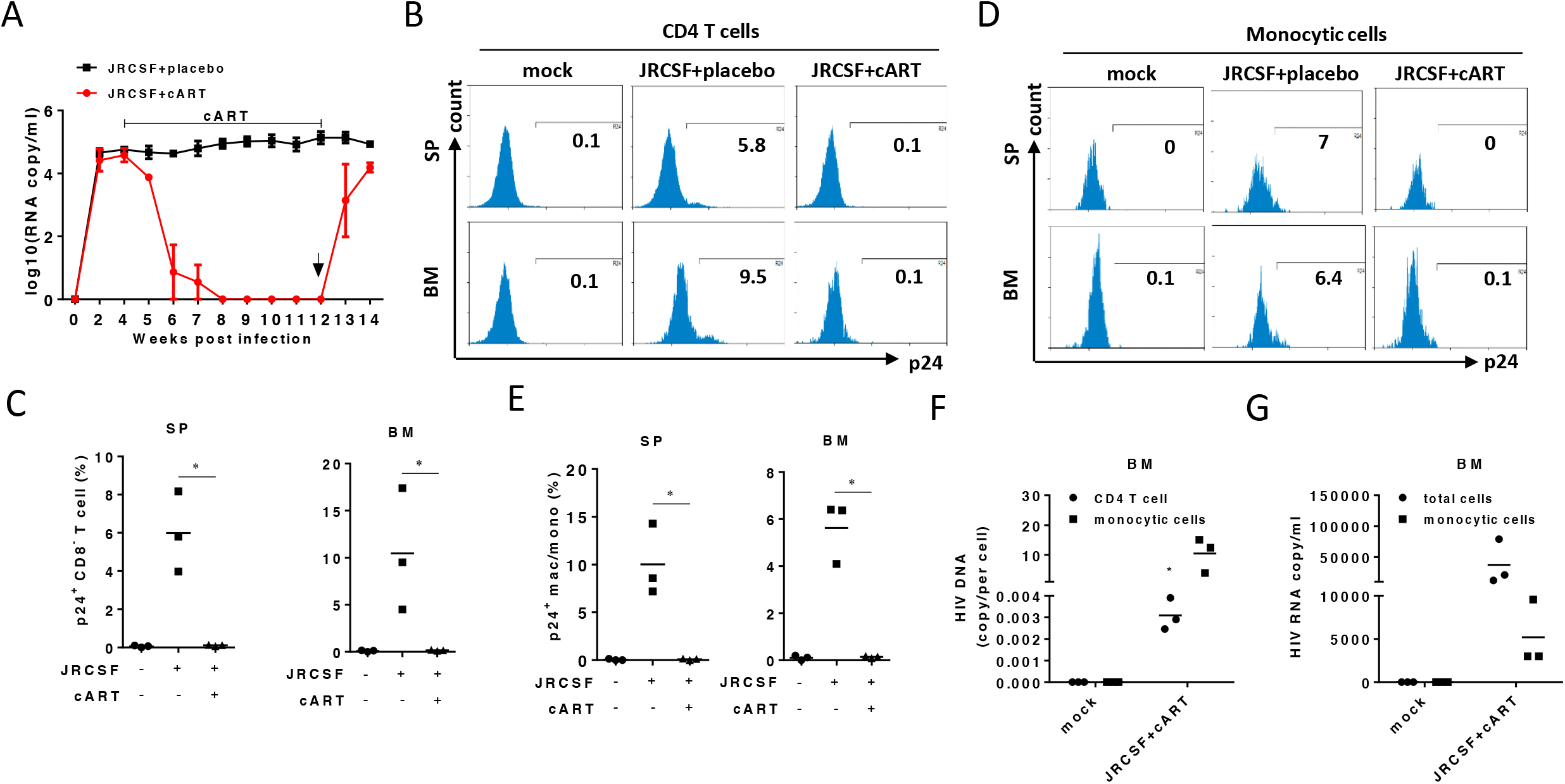
cART-resistant HIV-1 reservoirs in T cells and monocytic cells in humanized mice. Humanized mice were infected with HIV_JRCSF_. cART was initiated at 4 weeks after infection. Mice were terminated at 12 weeks after infection. Spleen and bone marrow cells were harvested for flow cytometry analysis. huCD45+CD3+CD8− (CD4 T cells including cells with HIV-induced CD4 downregulation) cells and huCD45+CD3-CD11c+CD14+ cells were purified by flow cytometer. Sorted cells were used for either PCR detection or virus out growth assay (VOA). (A) Plasma vial loads were measured weekly or every other week. (B) Histograms show percentage of HIV gag p24+ CD4 T cells (huCD45+CD3+CD8-) in the spleen and bone marrow. (C) Summarized percent p24+ cells of CD4+ (CD3+CD8-) T cells in the spleen and bone marrow. (D) Histograms show percent p24+ of monocytic cells (huCD45+CD3-CD11c+CD14+) in the spleen and bone marrow. (E) Summarized percent p24+ of monocytic cells in the spleen and bone marrow. (F) Relative HIV-1 DNA (copy per cell) in either CD4 T cells or monocytic cells. (G) Total bone marrow cells or purified monocytic cells were treated with TNF-α and co-cultured with MOLT-4 cells for 14 days. HIV-1 RNA in supernatants on day 14 were detected. Bars in dot graphs indicate mean value. Error bars indicate standard deviations (SD). * indicates p<0.05.

### Bromodomain inhibitor activates HIV-1 replication under suppressive cART in vivo

Bromodomain inhibitors have been shown to activate HIV-1 gene expression in different cell systems in vitro with latent or chronic HIV-1 infection (24-27). However, little is known about how bromodomain inhibitors affect HIV-1 reservoirs in vivo during cART. In order to investigate the effect of BETi on viral reservoirs in vivo, we first characterized the pharmacokinetic (PK) properties of I-BET151 in NSG mice. We showed that, at dose of 18mg/kg, the concentration of I-BET151 was maintained for at least 8 hours in blood and dropped below EC_50_ around 24 hours after the initial administration through gavage (Fig. S3). Thus, humanized mice were treated with I-BET151 once every 20 hours in the following experiments. To assess the capacity of I-BET151 in activating HIV-1 reservoir cells, HIV-1 infected hu-mice were treated with I-BET151 at 21wpi by daily gavage after viremia was completely suppressed for 4 weeks by cART. HIV-1 viremia rebound was detected on day 9 (22.3wpi) after I-BET151 treatment (Fig. 2A). To catch the putative HIV-1 reservoir cells responding to I-BET151 treatment, animals were terminated immediately after viremia rebound was detected. Leukocytes from spleens and bone marrow as well as tissues from liver and lung were harvested and subjected to the detection of cell-associated HIV-1 RNA by TaqMan real-time PCR. It demonstrated that blood viral load rebound was accompanied with a higher level of cell-associated HIV-1 RNA in human cells in different organs or tissues (Fig. 2B). However, there was no virus spreading or secondary infection occurred which is evidenced by a similar level of HIV-1 DNA in human cells in tissues upon the activation of viral replication after BETi treatment (Fig. 2C). Therefore, I-BET151 treatment activates HIV-1 gene expression, without spreading new infection, during cART in vivo.

**FIG 2.**
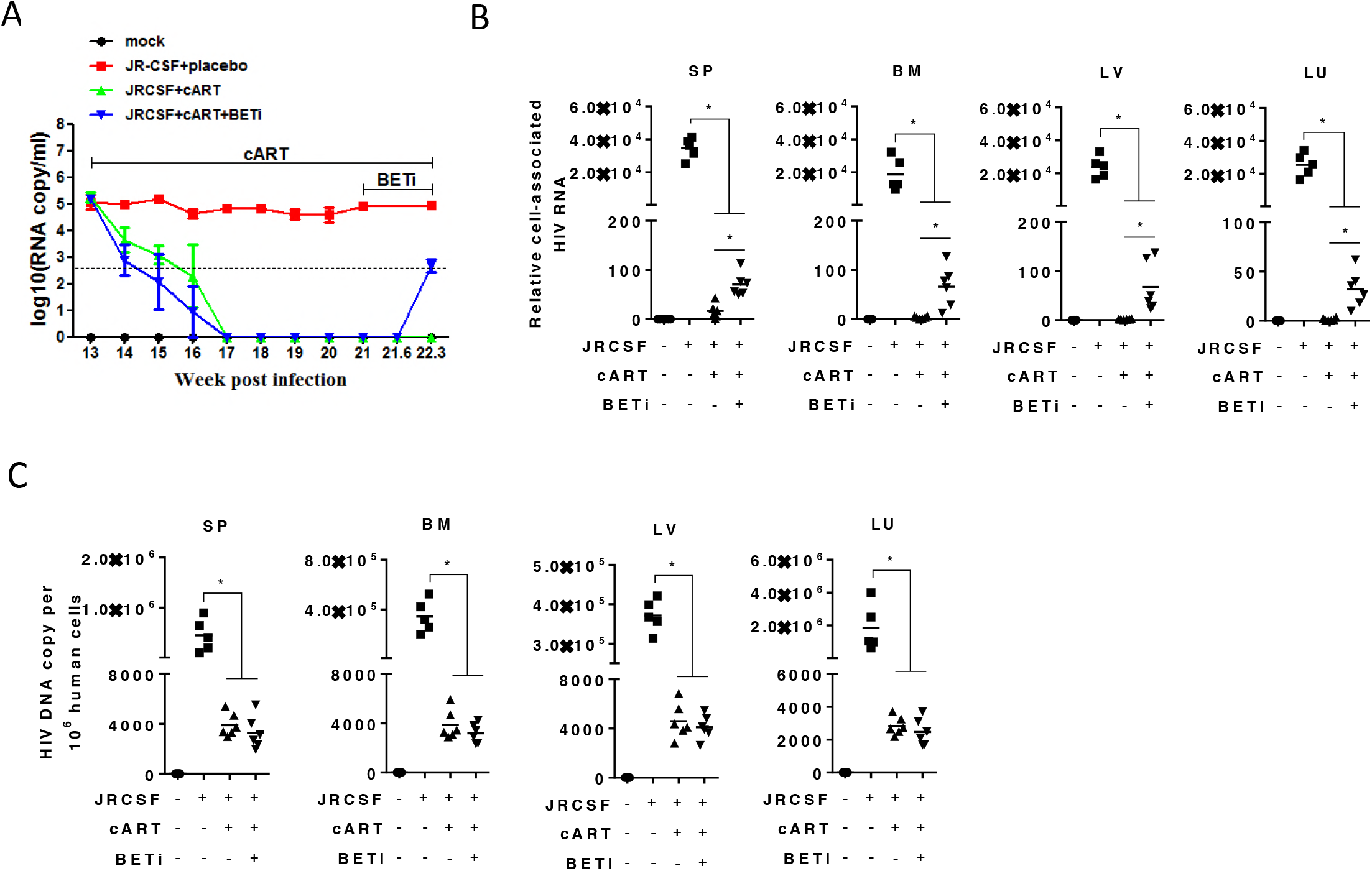
I-BET151 treatment activates HIV-1 replication under suppressive cART in vivo. Humanized mice infected with HIV_JR-CSF_ and were treated with cART initiated at 13 wpi. Mice were terminated at 22.3 wpi. (A) HIV-1 plasma viremia. (B) Relative cell-associated HIV-1 RNA levels in human cells or tissues (spleen/SP, bone marrow/BM, liver/LV and lung/LU), normalized to human CD4 mRNA levels. (C) Cell-associated HIV-1 DNA copy number in human cells in different tissues (spleen, bone marrow, liver and lung) was measured by real-time PCR. Bars in dot graphs indicate mean value. * indicate p<0.05.

### I-BET151 significantly activates HIV-1 replication in monocytic reservoir cells

In order to define the HIV-1 reservoir cells that were activated by BETi, we investigated what cell types were responding to I-BET151 treatment in vivo. Surprisingly, we did not observe any activation of p24 production in CD4 T cells (Fig. 3A and B), but a significant percentage of monocytic cells became p24-positive after I-BET151 treatment (Fig. 3C and D). And we did not observe significant p24 expression in other cell types (Data not shown). To confirm this finding, we performed immune-fluorescence staining on spleen tissue sections. Either human CD3/p24 or CD14/p24 were co-stained by specific antibodies. Consistently, we found p24 staining was co-localized with CD14 positive cells in I-BET151 group but not with CD3 positive cells. In comparison, no p24 positive cells were detected in cART-only mice, indicating an effective suppression of HIV-1 replication by cART (Fig. 3E). Accordingly, when we normalized cell-associated HIV-1 RNA to the level of human CD14 gene, a more significant increase of HIV-1 RNA level was observed as compare to normalization to human CD4 gene (Fig. 2B and Fig. S4). We calculated CD3^-^CD11c^+^CD14^+^ monocytic cell numbers in the spleen, and found I-BET151 did not alter the number of cells in treated group (Fig. 3F). However, the p24 positive monocytic cell number was significantly increased by I-BET151 treatment (Fig. 3G). Similar findings were also observed in the bone marrow (Fig. S5). In a separate experiment with sustained cART, the elevated viral load induced by I-BET15 treatment could be inhibited again when I-BET151 was stopped (Data not shown), indicating that the observed rebound of viral RNA production was not due to the emergence of drug-resistant mutant viruses. In addition, neither percentage nor total number of CD4^+^ T cells or huCD45^+^ cells in I-BET151-treated animals was affected in comparison to non-treated mice (Fig. S6). Thus, I-BET151 treatment did not mediate significant cytotoxicity in vivo in this study. Together, our results suggest that I-BET151 treatment preferentially activates HIV-1 replication in CD3^-^CD11c^+^CD14^+^ monocytic cells in comparison with CD4^+^ T cells under suppressive cART in vivo.

**FIG 3.**
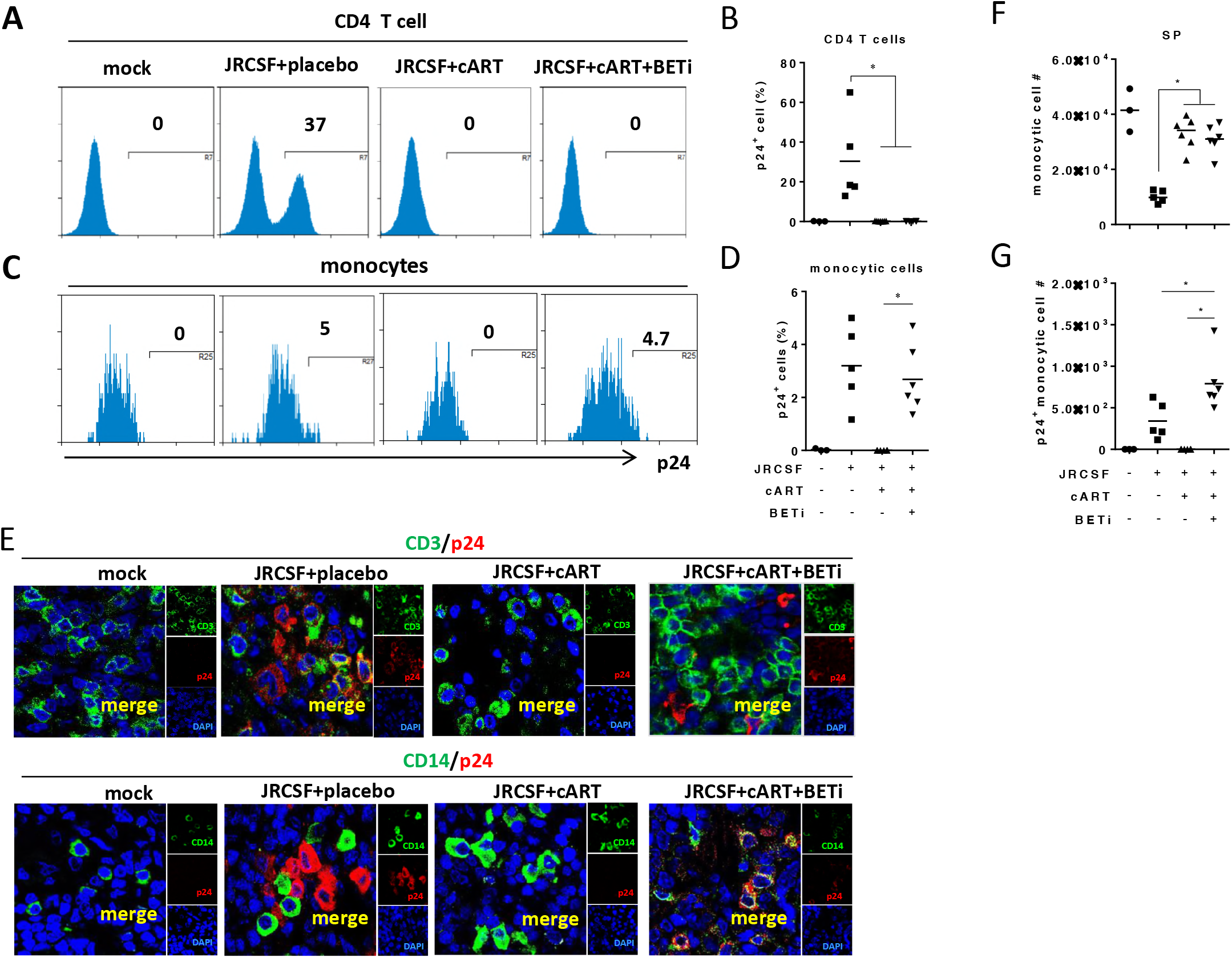
I-BET151 treatment specifically activates HIV-1 replication in monocytic cells under suppressive cART in vivo. Humanized mice were treated as in Fig. 2. (A) Histograms show percent HIV gag p24+ of CD4+ (CD3+CD8-) T cells in spleens. (B) Summarized percent p24+ of CD4+ T cells in spleens. (C) Histograms show percent p24+ of monocytc cells (CD3-CD11c+CD14+) in spleens. (D) Summarized percent p24+ of monocytc cells in spleens. (E) Immunofluorescence co-staining of CD3 (green)/gag p24 (red) (upper lane) or CD14 (green)/gag p24 (red) (lower lane) in spleens. (F) Monocytic cell numbers in spleens. (G) HIV gag p24+ monocytic cell numbers in spleens. Bars in dot graphs indicate mean value. * indicates p<0.05.

### I-BET151 activates HIV-1 reactivation more efficiently in monocytic cells than in T cells

It has been reported that BETi activates HIV-1 gene transcription in latent models of both T cells and monocytic cells in vitro (26, 27). To confirm our finding, we transduced either resting CD4^+^ T cells or monocyte derived macrophages (MDMs) with VSV-G HIV Duo-Fluo, which can distinguish productive infection from latent infection. This reporter virus encodes two separate fluorescent markers: an LTR-driven eGFP marker (productive infection) and an LTR-independent mCherry marker driven by an EF1α promoter (31, 32). After transduction, cells were cultured for 24 hours before I-BET151 treatment. mCherry and GFP expression was determined at 72 hours for CD4^+^ T cells and 36 hours for MDMs after culture with I-BET151(Fig. S7), at the concentration of 0.5μM at which I-BET151 induce minor cytotoxicity as tested on lymphocytic or monocytic cell lines (Fig. S8). We chose SAHA, an HDAC inhibitor reported to activate HIV LTR transcription in both cell lines and primary CD4 T cells (33-35), as positive control. The results showed that I-BET151 was unable to significantly enhance HIV-1 transcription in resting CD4^+^ T cells (Fig. 4A and B). In contrast, HIV-1 transcription in MDMs was significantly increased by I-BET151, as evidenced by the increase in the percent of GFP positive cells observed directly under fluorescence microscope, and confirmed by flow cytometry analysis (Fig. 4C-E).

**FIG 4.**
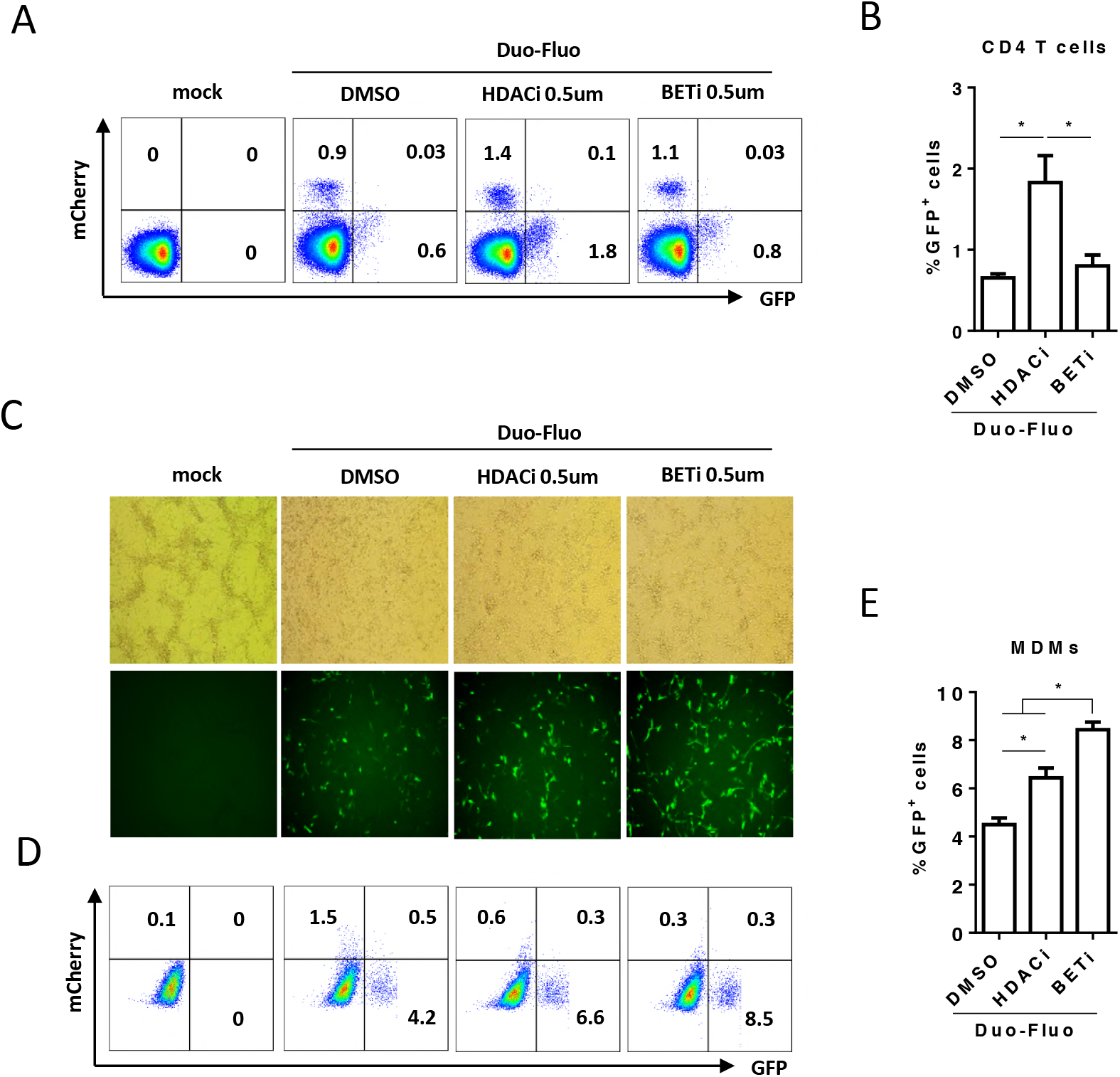
I-BET151 treatment activates HIV-1 transcription in MDMs but not in resting CD4+ T cells in vitro. CD25− resting CD4+ T cells and monocytes-derived-macrophages (MDMs) were transduced with VSV-G-G pseudotyped Duo-Fluo HIV-1 reporter virus and cultured for 24 hours before I-BET151 treatment. Cells were analyzed for mCherry and GFP expression at 72 hours (CD4+ T cells) or 36 hours (MDMs) after I-BET151 treatment. (A) Plots show mCherry and GFP expression in CD4+ T cells. (B) Summarized percentage of GFP+ cells in CD4+ T cells. (C) GFP frequency in MDMs was observed under fluorescence microscope. (D) Plots show mCherry and GFP expression in MDMs. (E) Summarized percentage of GFP+ cells in MDMs. Error bars indicate standard deviations (SD). * indicates p<0.05.

To further compare the relative efficiency of HIV-1 transcription activation by I-BET151 in T cells and monocytic cells, we performed experiments using lymphocytic clone (ACH-2) or monocytic clone (U1) with latent HIV-1 provirus. The titration results demonstrate that I-BET151 activated HIV-1 transcription in both ACH2 cells and U1 cells in a dose dependent manner. However, I-BET151 enhanced viral gene transcription in U1 cells at about 5-to-10 fold more efficiently than in ACH-2 cells using the same dose (Fig. 5A and 4B). An approximate 3-fold difference in EC_50_ was observed as calculated based on the viral transcription levels (Fig. 5C).

**FIG 5.**
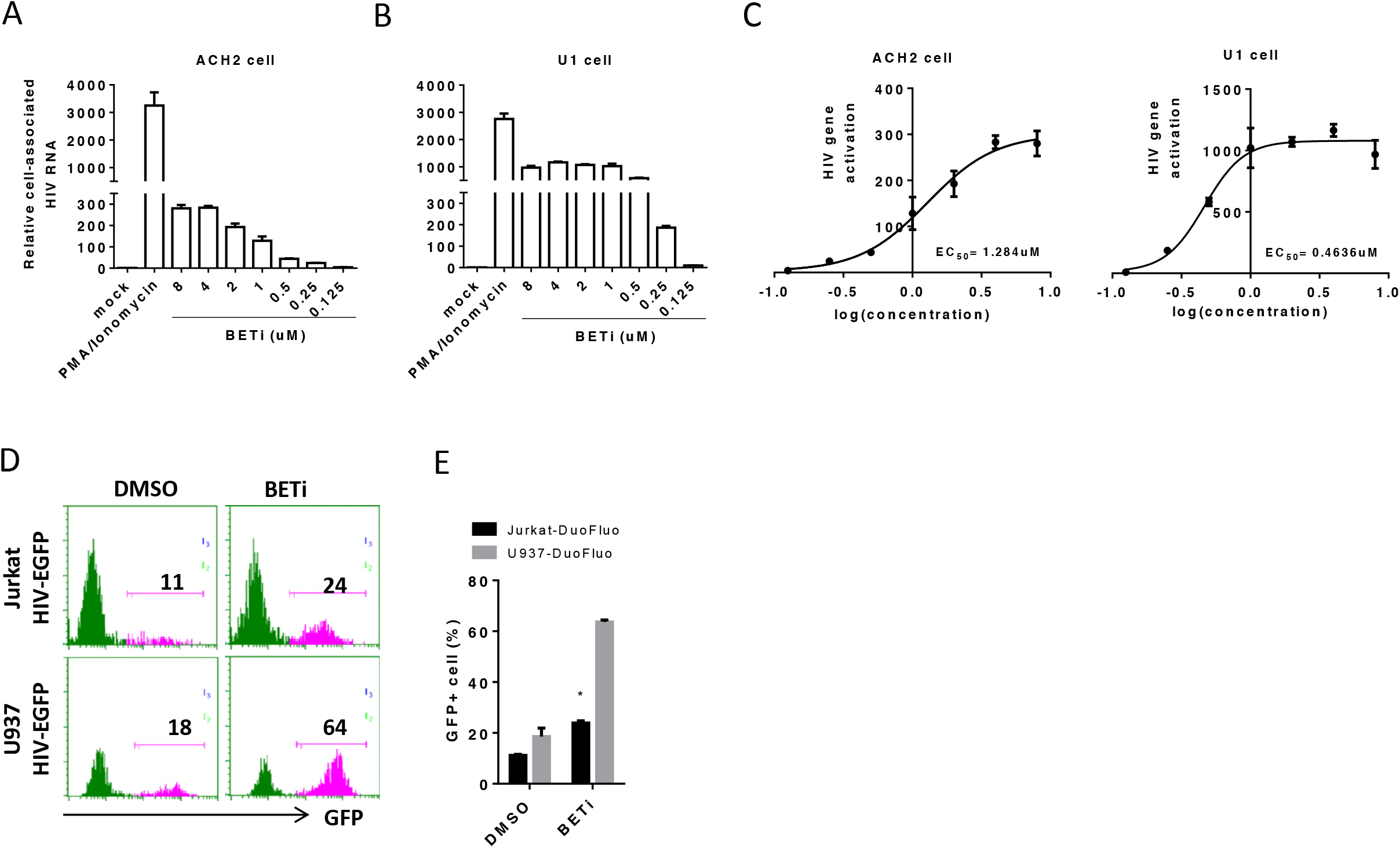
I-BET151 activates HIV-1 transcription in monocytic cells more efficiently than in lymphocytic T cells. ACH-1 T cells or U1 monocytic cells were treated with various concentrations of I-BET151. Cell-associated HIV gag RNA levels were detected by real-time PCR. (A) Relative viral RNA levels in ACH-2 cells at 48 hours post treatment. (B) Relative viral RNA levels in U1 cells at 48 hours post treatment. (C) EC50 of BET151 on ACH-2 or U1 cells calculated based on Fig. 4A or Fig. 4B. Jurkat cells or U937 cells were transduced with VSV-G-G pseudotyped Duo-Fluo HIV-1 reporter virus and mCherry positive only cells were purified by FACS. Sorted cells were expanded and treated with 0.5uM I-BET151 for 48 hours. (D) Percentages of GFP positive cells in Jurkat-DuoFuo (upper) or U937-DuoFluo (lower). (E) Summarized percentage of GFP+ cells in Fig. 4D. Error bars indicate standard deviations (SD). * indicates p<0.05.

We speculated that the quiescent status of HIV-1 provirus may contribute to this discrepancy observed in different cell types. Thus, we inoculated U937 cells and Jurkat cells with the dual reporter virus and sorted the mCherry^+^GFP^-^ cells (latent infection). After culturing for 4 days to expand the cells, we treated Jurkat-DuoFluo or U937-DuoFluo cells with I-BET151 (0.5μM) for 48 hours. The activation of HIV transcription was indicated by the percent of GFP positive cells after treatment. The results demonstrated that U937-DuoFluo was more sensitive to I-BET151 treatment as 64% cells became GFP positive comparing to 24% in Jurkat-DuoFluo (Fig. 5D and E). These results suggested that HIV-1 transcription in latent or low replicating monocytic cell reservoirs could be sufficiently activated or enhanced by I-BET151 as compared to latent T cell reservoirs.

### I-BET151 activates HIV-1 gene expression in monocytic cells via CDK2/CDK9-dependent mechanisms

Inhibition of either cyclin-dependent kinase (CDK) 2 or CDK9 has been reported to suppress HIV-1 transcription or replication in vitro or in vivo(36-41). Previous reports show that bromodomain inhibitors induced reactivation of HIV-1 latency is dependent on CDK9 (24, 25). HIV-1 replication, in CDK2-knockdown macrophages-like cells derived from pluripotent stem cells, was significantly reduced(42). To further study the mechanism of I-BET151-mediated HIV-1 transcription in monocytic cells and CD4^+^ T cells, we first established cell lines expressing luciferase under the control of HIV-1 LTR using either U937 (U937-luc) or Jurkat (Jurkat-luc) cells. We treated cells with I-BET151 in the presence of either CDK2 inhibitor (K03861)(43) or CDK9 inhibitor (LDC000067), and cultured for 48 hours before luciferase activity detection. In U937-luc cells, either CDK2 or CDK9 inhibitor alone could down-regulate HIV-1 transcription relative to mock cells. When I-BET151 was introduced, the enhancement of HIV-1 transcription was abolished by either CDK2 (1μM) or CDK9 (25μM) inhibitors (Fig. 6A). Interestingly, CDK2 inhibitor failed to inhibit I-BET151-induced HIV-1 transcription, even increased HIV-1 transcription in Jurkat-luc cells. While CDK9 inhibitor was able to down-regulate HIV-1 transcription and inhibit the activity of I-BET151 to enhance HIV transcription in Jurkat-luc cells (Fig. 6B). These results suggested that the different efficacy of I-BET151 in activating HIV-1 transcription in monocytic cells and CD4^+^ T cells is attributed to cell-intrinsic difference in the regulation of HIV-1 transcription by CDK2.

**FIG 6.**
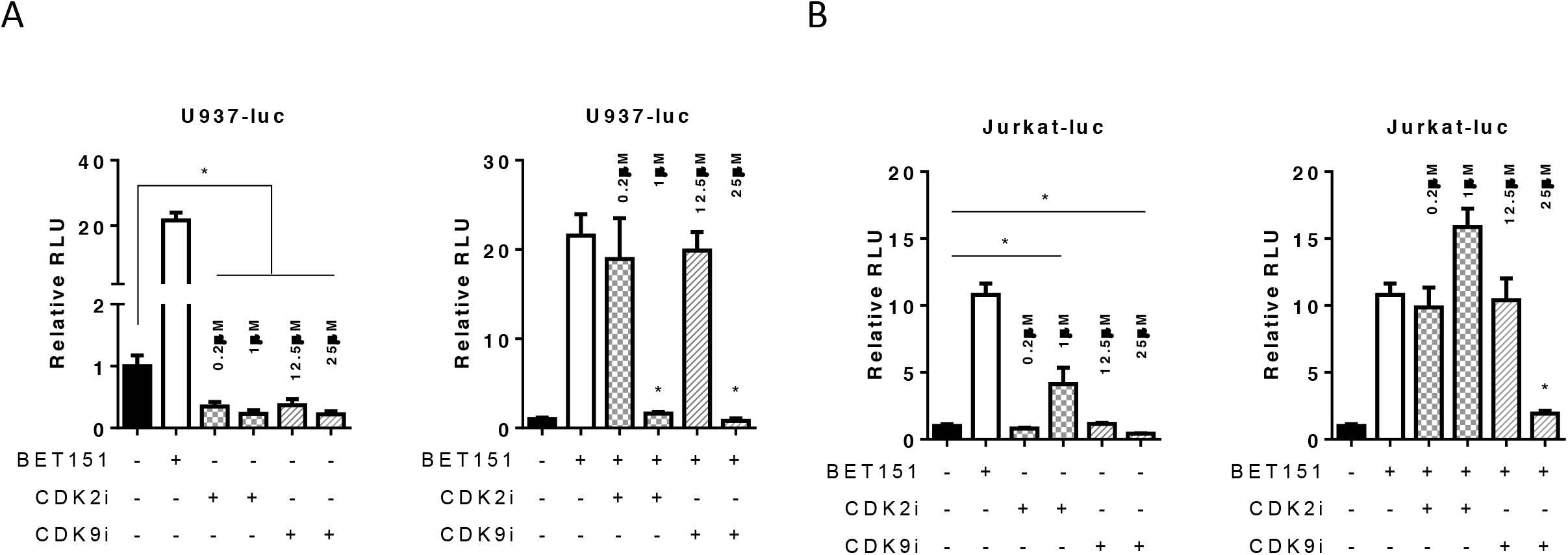
I-BET151-activated HIV replication in monocytic cells is dependent on both CDK2 and CDK9. U937 cells and Jurkat cells were transduced with VSV-G-G pseudotyped HIV-luc reporter virus. After expansion for 6 days, cells were treated with CDK2 or CDK9 inhibitor with or without I-BET151 treatment. At 48 hours after I-BET151 treatment, luciferase activity in cells were measured. (A) Luciferase activity in U937 cells after CDK2 or CDK9 inhibitor treatment with or without I-BET151. (B) Luciferase activity in Jurkat cells after CDK2 or CDK9 inhibitor treatment with or without I-BET151. Error bars indicate standard deviations (SD). * indicates p<0.05.

## DISCUSSION

Despite the importance of HIV-1 reservoirs in resting CD4^+^ T cells, it is impossible to achieve a cure without considering reservoirs cells like macrophages in brain tissues (44, 45). The existence of HIV-1 macrophages reservoirs in vivo is strongly supported by Honeycutt et al (17), who demonstrated that HIV-1 replication can be sustained in macrophages in a humanized mice model without T cells. Consistently, our study demonstrates that monocytes or macrophages are important HIV-1 reservoir cells during suppressive cART. Importantly, bromodomain inhibitors could be a novel measure adopted to activate viral replication in this particular reservoir in vivo.

It is known that HIV-1 reservoir cells can be exit in two different statuses in vivo. One is harboring latent replication-competent proviruses expressing little to no viral products. The long-lived resting memory CD4^+^ T cells is the putative latent reservoir cells mostly studied to date (46-48). The other kind of reservoir cells is the residual active reservoir cells which consistently produce low level of viruses, because of either sanctuary sites that have low accessibility to cART, or the cell types that less responsive to certain anti-retroviral drugs such as monocytes or macrophages(49-51). The very recent report proved the existence of lymphoid tissue sanctuary sites for harboring low level of viral replication in patients with undetectable viremia (52). And the brain resident macrophages have been considered to play a pivotal role in maintaining HIV-1 persistent infection since cART in brain is not as effective as in other tissues (44, 45). In our study, we cannot distinguish if the increased level of HIV-1 replication in monocytic cells is due to activation of HIV provirus or enhancement of residual low level of viral replication. However, our findings are valuable for HIV reservoir study and development of strategies targeting to different persistently infected cells during cART.

In this study, no reduction of cell-associated HIV-1 DNA in human cells was observed solely by I-BET151 treatment on day 9 after the administration. We speculate that the poor elimination of HIV-1 reservoir induced by I-BET151 in vivo is due to two reasons. First, macrophages are more resistant to HIV-1 infection induced cytopathic effect comparing to CD4^+^ T cells (53, 54). As I-BET151 effectively reactivates HIV-1 replication only in monocytic cells in vivo, it is likely that the cells are simply not dying while producing viruses. Second, the immune system might be still defective or exhausted due to chronic HIV-1 infection though viral replication being suppressed by cART. Given the above possibilities, bromodomain inhibitor treatment should be joint with therapy can either kill HIV-1 positive cells directly (immune-toxin) or recover immune response to HIV-1 infection(29). Thus, bromodomain inhibitor therapy requires more interventions to synergistically purge HIV-1 reservoir cells.

P-TEFb is composed of Cyclin T1 and CDK9, and converts promoter-proximally paused RNA polymerase II (RNAPII) complexes into efficient elongating complexes(23). In the absence of HIV-1 trans-activator Tat, RNAPII pauses after transcription of the TAR sequence. Tat recruits P-TEFb via CycT1 to TAR, allowing CDK9 to phosphorylate the C-terminal domain (CTD) of RNAPII and continues viral gene transcription thereafter (55). It has been reported that CDK2 can phosphorylate either CDK9 or Tat, and thereby contributes to enhanced HIV-1 transcription(56, 57). These findings suggest both CDK9 and CDK2 are involved in the regulation of HIV-1 transcription. However, CDK2 expression is low or undetectable in resting CD4^+^ T lymphocytes(58), which is likely to be associated with the failure of CDK2 inhibitor to exert suppression on viral transcription with or without I-BET151 treatment in our study. To further understand the role of CDK2 in the regulation of HIV-1 gene expression and latency in monocytic cells will be beneficial for future HIV-1 cure study.

Although the mechanisms contributing to HIV-1 transcription quiescence have investigated extensively, we still do not fully understand the relative importance of each mechanism in different cell types. As we have recently reported, depletion of regulatory T cells during cART in humanized mice leads to reactivation of HIV-1 replication predominantly in memory CD4^+^ T cells(28). In addition, it is of particular importance to identify all possible HIV-1 reservoir cells in lymphoid organs in vivo. To investigate how HIV-1 maintains its low replication or latency in different cell type will help guide future drug design.

## ACKNOWLEDGMENTS

We thank Qiong He and Adrain Kingsberry for technical assistance; We thank the University of North Carolina (UNC) Division of Laboratory Medicine for animal care; the UNC CFAR, Flow Cytometry Core Facility, and the UNC Animal Histopathology Core Facility. We thank all members in the Su laboratory for critical reading and/or discussion of the paper, and for their input and assistance. This study was supported in part by UNC University Cancer Research Fund innovation grant, and the US National Institutes of Health (AI080432, AI077454 and AI095097 to LS). This work was also supported by GSK. GSK participated in study design.

## MATERIALS AND METHODS

### Construction of humanized mice

Human fetal liver tissues were obtained from elective or medically indicated termination of pregnancy through a non-profit intermediary working with outpatient clinics (Advanced Bioscience Resources). Approval for animal work was obtained from University of North Carolina (UNC) Institutional Animal Care and Use Committee (IACUC). We constructed hu-NRG mice as previously reported (28, 59-63).

### HIV-1 infection of humanized mice

Humanized mice were infected by intravenous (i.v.) injection with HIV-1_JRCSF_ stocks (10 ng p24/mouse in 50μl) or with mock stocks in control mice.

### cART regimens in NRG-hu HSC mice

Individual tablets of TRUVADA (tenofovir/emtricitabine; Gilead Sciences) or raltegravir (Merck) were crushed into fine powder and manufactured as 5BXL by TestDiet based on previously published (29, 64).

### Luciferase assay

Cells were transduced with VSV-G HIV-luc, in which luciferase gene expression is under control by HIV LTR, at dose of 1 MOI. After culture for 6 days, cells were treated with CDK2 or CDK9 inhibitor alone, or in the presence of I-BET151. At 48 hours after treatment, cells were washed with PBS and lysed in 100μl passive lysis buffer (Promega). Cell lysis were transferred into 96 well plates and luciferase activity was detected by adding 50 μl substrate into each well and read by 96 microplate luminometer (GLOMAX).

### Cell sorting and culture

For primary cells, bone marrow cells from humanized mice were harvested and submit to cell sorting by flow cytometer (ARIA II, BD). Human CD45^+^CD3^+^CD8^-^ cells and CD45^+^CD3^-^ CD11c^+^CD14^+^ cells were sorted. For cell lines, U937 and Jurkat cells were first transduced with VSV-G Duo-Fluo reporter virus at dose of 1 MOI and expand for 6 days. GFP-mCherry+ cells were sorted by ARIA II (BD) and expand for 6 days in 10% FBS RPMI-1640 before experiment. The purity of sorted cells was all over 99% (Data not shown).

### Quantification of HIV viral load in plasma

Peripheral blood was collected with EDTA as an anti-coagulant at indicated time points after HIV infection. Plasma was prepared by centrifugation and stored at -80°C until assay. Viral RNA was isolated from the plasma. HIV viral road was measured by qRT-PCR as described previously(28, 29, 60).

### Cell-associated HIV-1 DNA

Total nucleic acid was extracted from cells or tissues using DNeasy mini kit (Qiagen). HIV-1 DNA was quantified by real-time PCR. Genomic DNA of ACH2, which contains one copy of HIV genome in each cell, was serially diluted in mouse leukocytes DNA to generate a standard curve(29).

### Cell-associated HIV-1 RNA

Total RNA was extracted from cells or tissues using RNeasy plus mini kit (Qiagen). HIV-1 RNA was detected as previously described(28, 29, 60). The HIV-1 gag RNA expression were normalized to human CD4 or CD14 mRNA level and relative HIV-1 gene expression levels were calculated according to 2^-ΔΔCT^ (29, 65).

### Cell lines and culture

ACH-2, Jurkat, U-937and U1 cells were obtained through the National Institutes of Health (NIH) AIDS Research and Reference Reagent Program, Division of AIDS. Cells were grown in RPMI 1640 (Gibco) with 10% fetal bovine serum (FBS) (Invitrogen), 5% penicillin-streptomycin (Invitrogen), and 2 mM glutamine (Invitrogen). Cells were maintained at a concentration of 10^6^ cells/ml in T-175 flasks. Cell concentrations and cell viability were monitored throughout the experiment at all time points studied.

### Virus outgrowth assay

Splenocytes of humanized mice (1x10^6^, 2x10^5^, 4x10^4^ human cells in single, duplicates or triplicates) were stimulated with PHA (2μg/ml) and IL-2 (100 units/ml) for 24 hours. MOLT4/CCR5 cells were added on day 2 to enhance the survival of leukocytes and to support HIV-1 replication. Culture medium containing IL-2 and T cell growth factor (29) was replaced on days 5 and 9. After 7 and 14 days of culture, supernatant was harvested individually and HIV-1 RT-qPCR was performed to score viral outgrowth. Estimated frequencies of cells with replication-competent HIV-1 were calculated using limiting dilution analysis.

For sorted cells, five hundred of each sorted cells were cultured in the presence of TNF-α (5ng/ml) in 100ul RPMI-1640 medium complemented with 10% FBS for 24 hours in U bottom 96-well plate and then 5x10^4^ MOLT4/CCR5 cells in 100ul same medium were added into each well, and co-culture for 14 days with half medium change every 3 days. 150ul supernatant were used for measuring viral load by TaqMan real-time PCR.

### Primary cell isolation

Human blood buffy coats were ordered from Gulf Coast Regional Blood Center and PBMCs were separated with Ficoll (Ficoll-Paque™ PLUS, GE Healthcare). Resting CD4^+^ T cells were purified by negative selection using a CD4 separation kit (Miltenyi Biotec) according to the manufacturer’s instructions. CD25^-^CD4^+^ T cells were subsequently isolated from CD4^+^ T cells by negative selection using CD25 beads (Miltenyi Biotec). CD14+CD16− monocytes were isolated using the EasySep- Human Monocyte Isolation Kit (STEMCELL). All cell purifications were performed according to manufactures’ instruction. Purity of CD25^-^CD4^+^ T cells was checked by stain with anti-CD4-FTIC and anti–CD25-PE (Biolegend) Purity of monocytes was checked by stain with anti-CD3-FITC and anti-CD14-PE (Biolegend). Cell purity was analyzed by Guava Easycyte™ 8HT (EMD Millipore). The purity of each isolated population was over 90%(data not shown).

### Primary cell transduction and culture

CD4^+^ T cells were transduced with VSV-G Duo-Fluo reporter virus at 0.1 MOI and cultured in RPMI-1640 complemented with 10% FBS and IL-2 (10u/ml) for 24 hours before I-BET151 treatment. Monocytes-derived macrophages (MDMs) were obtained by culturing monocytes with M-CSF (50ng/ml) and GM-CSF (50ng/ml) in RPMI-1640 with 10% FBS for 7 days. MDMs were harvested and transduced with VSV-G Duo-Fluo reporter virus at 0.1 MOI and cultured for 24 hours before I-BET151 treatment.

### Inhibitors and treatment

Bromodomain inhibitor I-BET151 was obtained from GlaxoSmithKline (Research Triangle Park, NC). Drug powder was dissolved in 0.5%HPMC:0.1%Tween80 to a final concentration of 1.5 mg/ml and pH 5. Administration of the drug was through daily oral gavage at dose of 18mg/kg. The solvent without drug was used as placebo. Highly potent and specific CDK2 inhibitor (K03861) and CDK9 inhibitor (LDC000067) (Selleckchem) were dissolved in DMSO and diluted to the indicated concentration in medium before use.

### Flow cytometry

For HIV-1 gag p24 staining, cells were stained with surface markers first, and then permeabilized with cytofix/cytoperm buffer (BD Bioscience, cat#554714), followed by intracellular staining. PE-conjugated anti-human CD11c (clone:3.9, cat#301606), PE/Cy5-conjugated anti-human CD4 (clone:RP4-T4, cat#300510), PE/Cy7-conjugated antihuman CD3 (clone:HIT3a, cat#300316), Pacific blue-conjugated anti-human CD14 (clone:HCD14, cat#325616), and APC/Cy7-conjugated anti-human CD45 (clone:H130, cat#304014) were purchased from Biolegend; Pacific orange–conjugated anti–mouse CD45 (clone:HI30, cat#MHCD4530), PE-TR-conjugated CD8 (clone:3B5, cat#MHCD0817), and LIVE/DEAD Fixable Aqua Dead Cell Stain Kit (cat#L34957) were purchased from Invitrogen. FITC-conjugated anti-HIV p24 (clone: FH190-1-1, cat#6604665) was purchased from Beckman Coulter. Flow cytometry was performed using either CyAn ADP (DAKO) or BD LSRFortessa (BD Biosciences), analyzed by Summit 4.3 (Beckman Coulter) or FlowJo 10 (FLOWJO, LLC), accordingly.

### Immunofluorescence staining of spleens

Mice spleens were harvested, fixed with 10% formalin (Fisher, Fair Lawn, NJ), embedded in paraffin and cut into 5μm tissue sections. Antigen retrieval was performed by incubation in Diva Decloacker (Biocare Medical, Concord, CA) for 30 min at 95°C, followed by slow cooling down for one hour. The tissue section was blocked with Background Sniper (Biocare Medical, Concord, CA). The sections were then stained with the primary antibodies: rabbit monoclonal anti-human CD3 (Life Span Bio Sciences, Seattle, WA; 1:100 dilution) or rabbit monoclonal anti-human CD14 (Abcam; 1:500 dilution) or mouse monoclonal anti-HIV-1 p24 (Dako, 1:5 dilution) diluted in blocking buffer (PBS, 0.05% Tween 20, 5% goat serum), and then secondary antibodies: Alexa Fluor 594 Donkey Anti-Mouse IgG (Life Technologies, Eugene, OR) and Alexa Fluor 488 Donkey Anti-Rabbit IgG (Life Technologies, Eugene, OR). The sections were stained with DAPI and mounted with an anti-fade mounting media (Abcam Cambridge, MA). Sections were analyzed by confocal microscopy (Zeiss LSM 700 Confocal Laser Scanning Microscope).

### Statistical analysis

Unpaired 2-tailed Student t tests and OneWay Anova analysis of variance with the Bonferroni multiple comparison test were performed using GraphPad Prism (GraphPad Software, San Diego, CA). P value of < 0.05 was considered statistically significant. All data were reported as mean ± SD (Standard deviation).

## Highlights

- BET-151 preferentially activate HIV-1 infection in CD3-CD11c+CD14+ monocytic cells as compare to CD4 T cells during suppressive cART in vivo.
- CD3-CD11c+CD14+ monocytic cells harbor replication-competent provirus during cART.
- BET-151 reactivate HIV-1 transcription and enhance viral replication in CD3-CD11c+CD14+ monocytic cells more than in resting CD4 T cells.
- BET151-induced activation of HIV-1 replication in monocytic cells is dependent on both CDK2 and CDK9, whereas only CDK9 is involved in the activation of HIV-1 replication mediated by BET151 in T cells.

---

**FIG S1.**
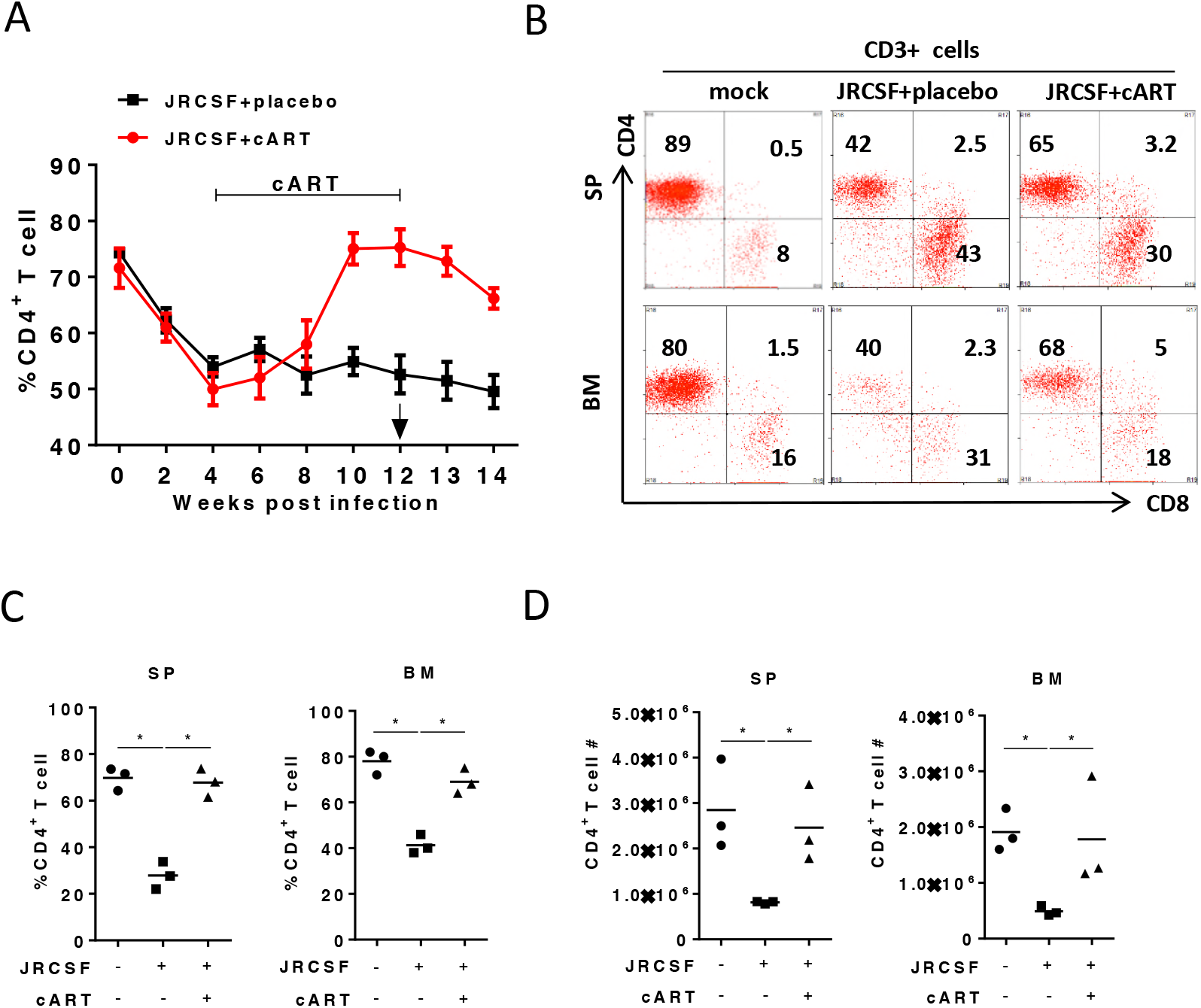
Humanized mice were treated and terminated as in Fig. 1A. (A) CD4+ T cell percentages in CD3+ cells over time in peripheral blood. (B) Representative plots show percentage of CD4+ T cells in CD3+ cells in spleen and bone marrow. (C) Summarized percentage of CD4+ T cells in CD3+ cells in spleen and bone marrow. (D) CD4+ T cell numbers in spleen and bone marrow. Bars in dot graphs indicate mean value. * indicate p<0.05

**FIG S2.**
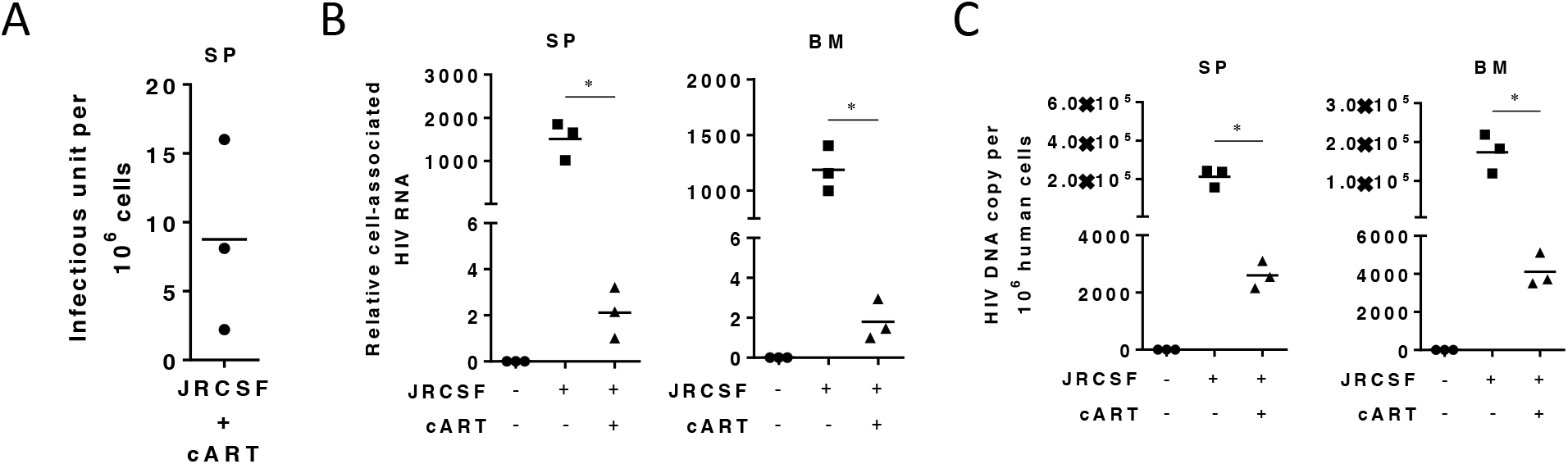
cART-resistant viral reservoir persist during suppressive cART in humanized mice. Humanized mice were treated and terminated as in Fig. 1A. (A) Replication-competent HIV-1 reservoir cells detected by virus outgrowth assay. (B) Relative cell-associated viral RNA in cells from spleen and bone marrow. (C) Cell-associated viral DNA in cells from spleen and bone marrow. Bars in dot graphs indicate mean value. * indicate p<0.05.

**FIG S3.**
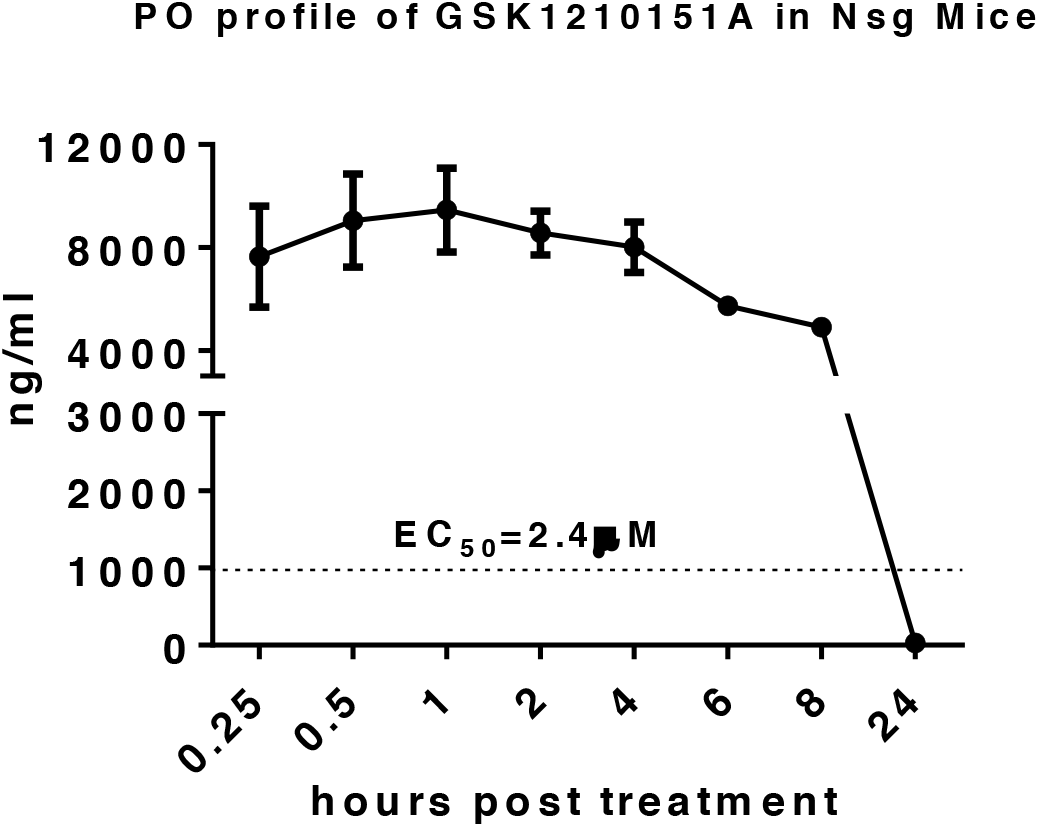
Pharmacokinetics of I-BET151 in NSG mice. NSG mice were treated with 18mg/kg I-BET151 through gavage and drug concentration in blood was measured by HPLC.

**FIG S4.**
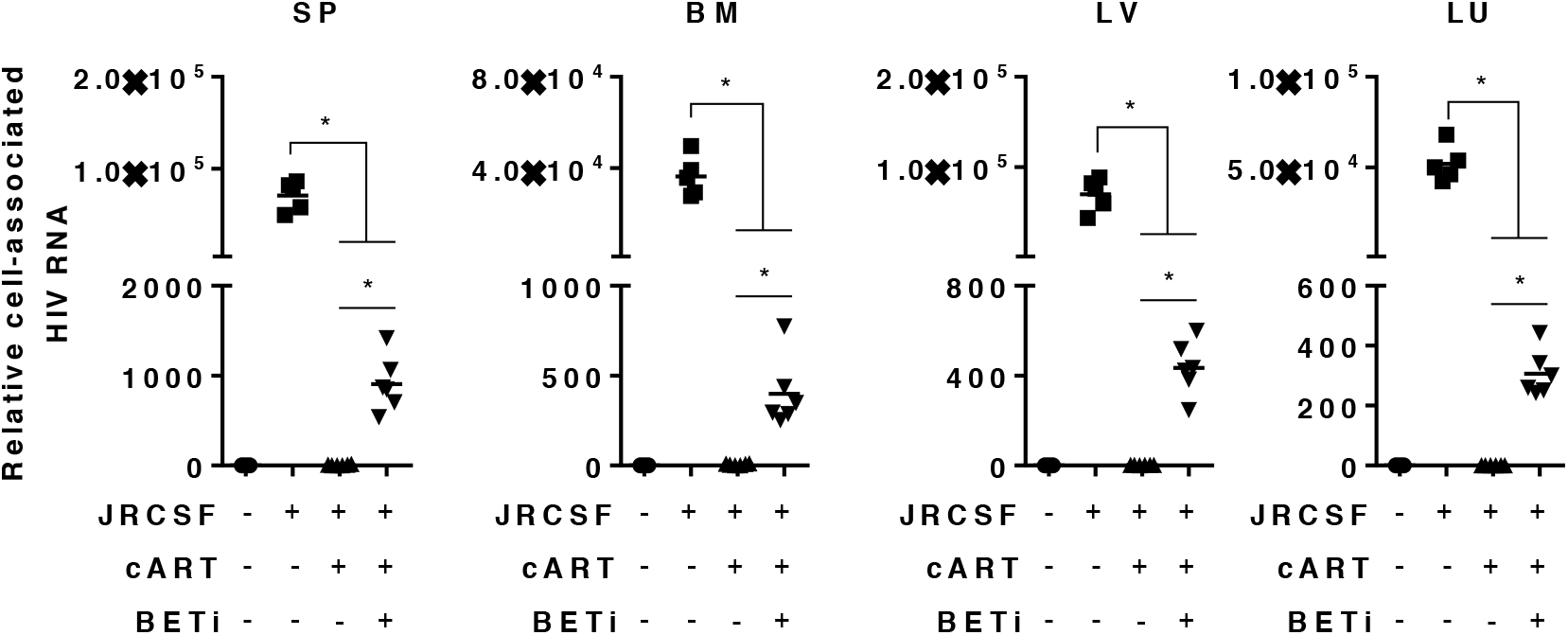
I-BET151 treatment activates HIV-1 replication under suppressive cART in vivo. Relative cell-associated viral RNA levels in human cells or tissues, normalized to human CD14 mRNA level. Bars in dot graphs indicate mean value. * indicate p<0.05.

**FIG S5.**
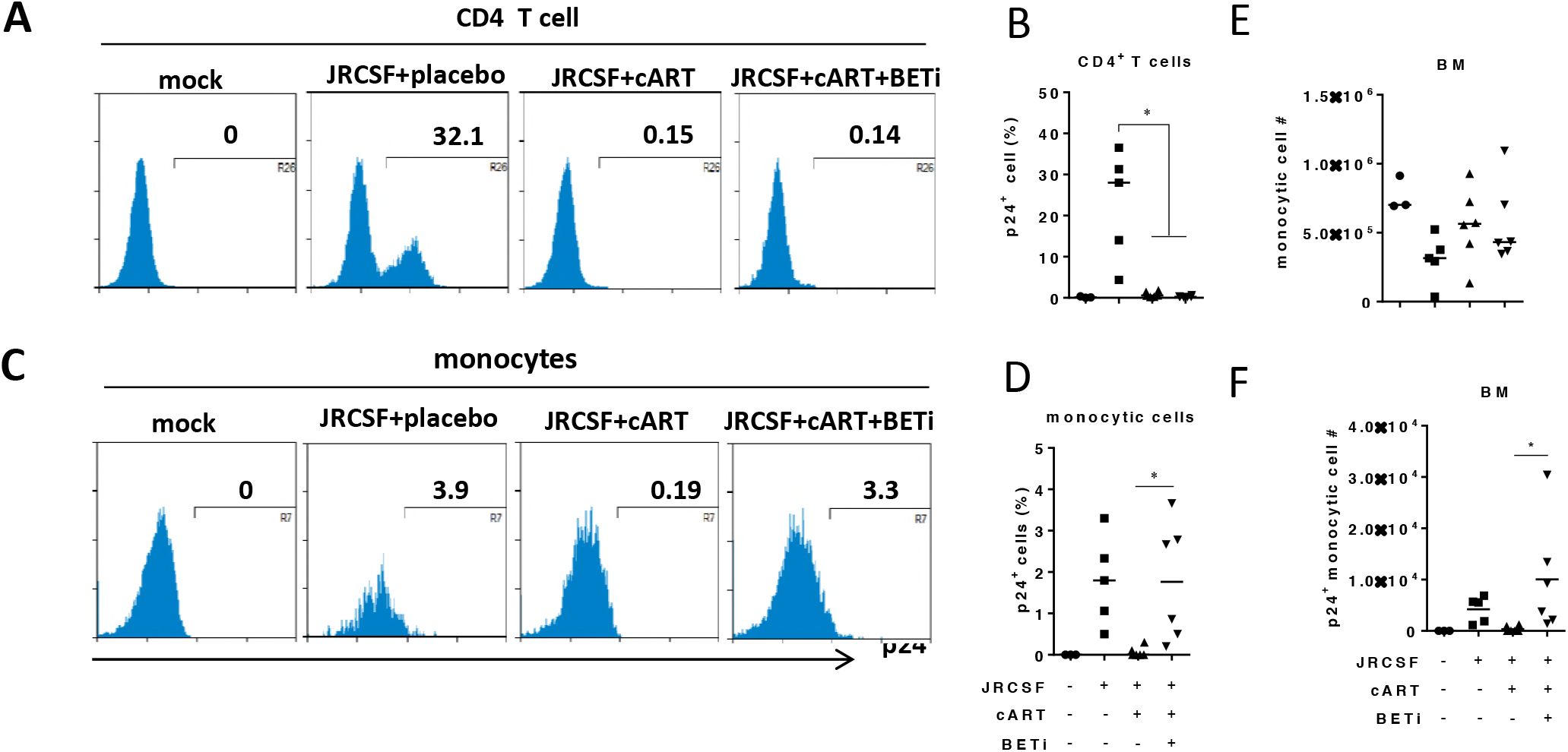
I-BET151 treatment preferentially activate HIV replication in monocytes under suppressive cART in bone marrow. Humanized mice were treated as in Fig.2 and bone marrow cells were investigated. (A) Histograms show percentage of HIV gag p24+CD4+ T cell (CD3+CD8-). (B) Summarized percentage of p24+CD4+ T cell. (C) Histograms show percentage of HIV gag p24+monocytc cells (CD3-CD11c+CD14+). (D) Summarized percentage of p24+monocytc cells. (E) Monocytic cell numbers. (F) HIV gag p24+ monocytic cell numbers. Bars in dot graphs indicate mean value. * indicate p<0.05.

**FIG S6.**
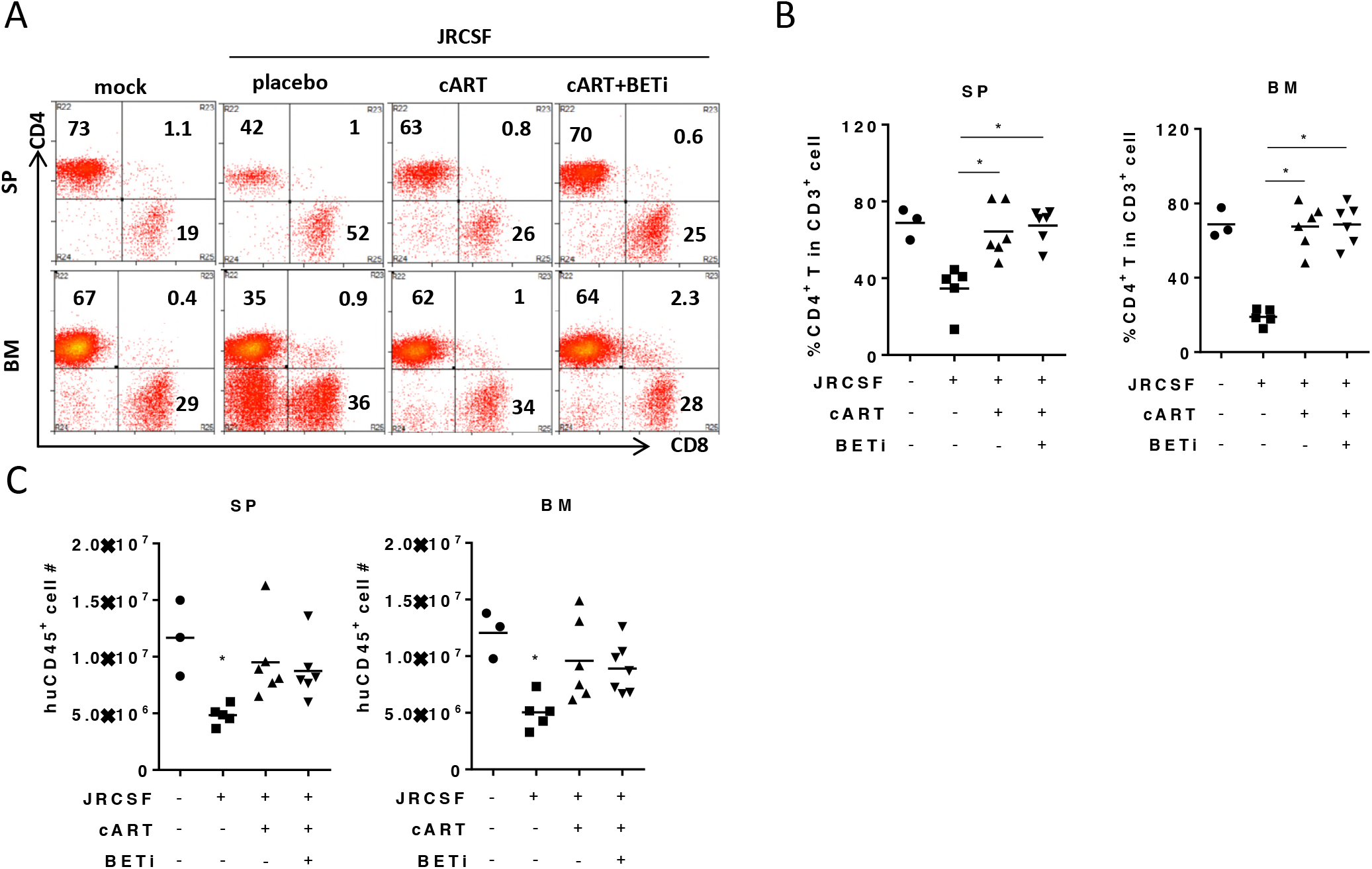
I-BET151 treatment does not affect T cells and total human leukocytes in vivo. On termination as in Fig.2, cells were harvested from spleen or bone marrow for flow cytometry analysis. (A) Plots show percentages of CD4+ T cell and CD8 T cell in CD3+ cells in spleen and bone marrow. (D) Summary data show percentages of CD4+ T cell and CD8 T cell in CD3+ cells in spleen and bone marrow. (C) Human CD45+ cells numbers in spleen and bone marrow. Bars in dot graphs indicate mean value. * indicate p<0.05.

**FIG S7.**
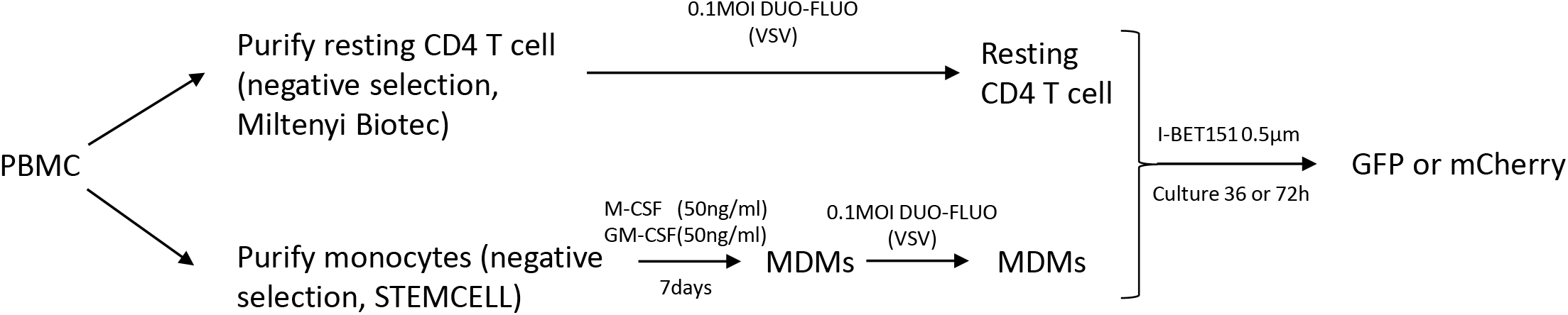
Flow chart for ex vivo assay

**FIG S8.**
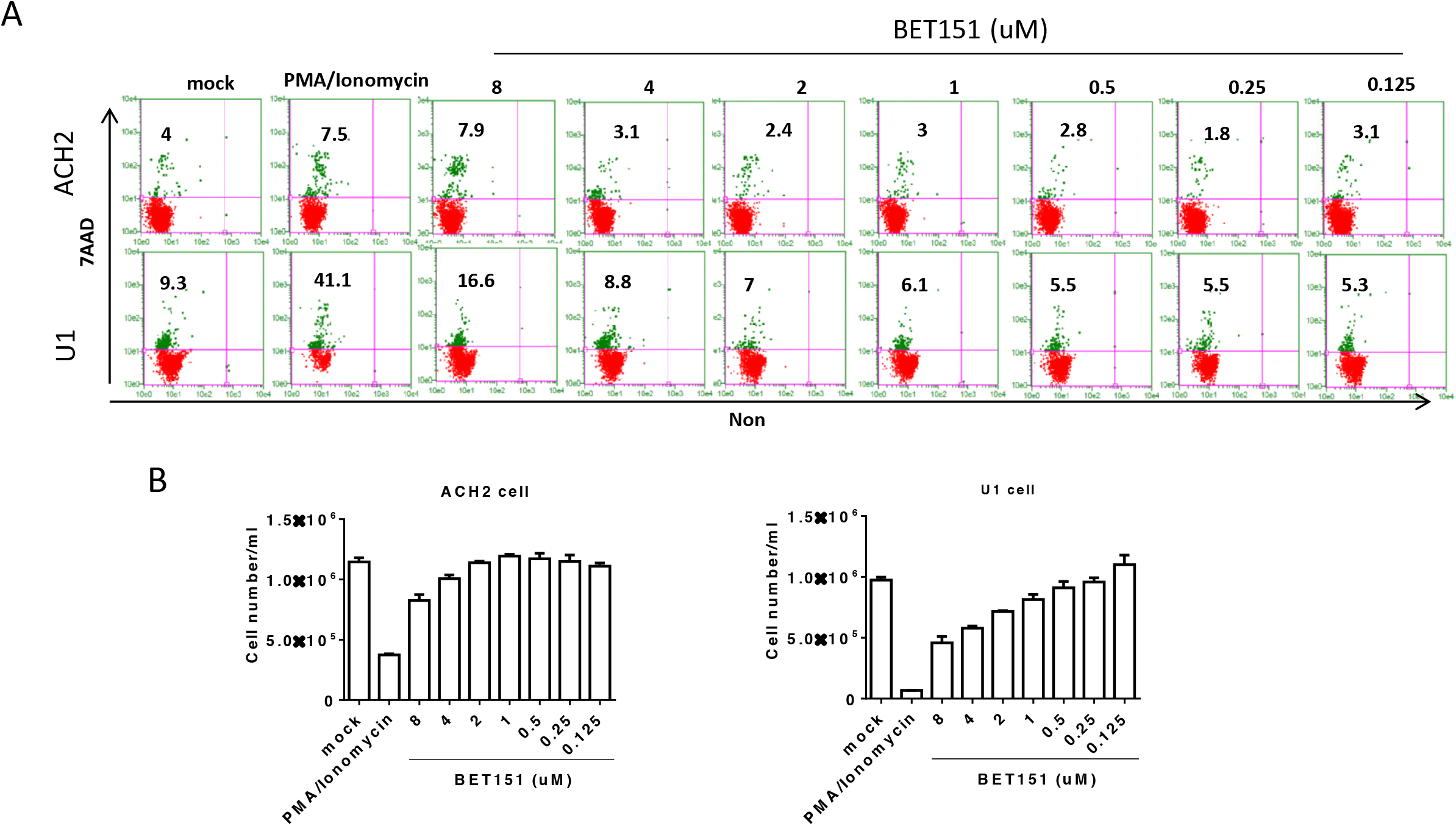
I-BET151 treatment induced cytotoxicity. ACH-1 or U1 cells were treated with series concentration of I-BET151 in cell culture. (A) Cell viability as evidenced by 7-AAD positive cells. (B) Live cell numbers were counted by Guava Pro system.

## REFERENCES

1. Perelson AS, Essunger P, Cao Y, Vesanen M, Hurley A, Saksela K, Markowitz M, Ho DD. 1997. Decay characteristics of HIV-1-infected compartments during combination therapy. Nature 387:188–191.

2. Palella FJ, Jr., Delaney KM, Moorman AC, Loveless MO, Fuhrer J, Satten GA, Aschman DJ, Holmberg SD. 1998. Declining morbidity and mortality among patients with advanced human immunodeficiency virus infection. HIV Outpatient Study Investigators. The New England journal of medicine 338:853–860.

3. Broder S. 2010. The development of antiretroviral therapy and its impact on the HIV-1/AIDS pandemic. Antiviral research 85:1–18.

4. Hsu DC, Sereti I. 2016. Serious Non-AIDS Events: Therapeutic Targets of Immune Activation and Chronic Inflammation in HIV Infection. Drugs 76:533–549.

5. Cihlar T, Fordyce M. 2016. Current status and prospects of HIV treatment. Current opinion in virology 18:50–56.

6. Suphanchaimat R, Sommanustweechai A, Khitdee C, Thaichinda C, Kantamaturapoj K, Leelahavarong P, Jumriangrit P, Topothai T, Wisaijohn T, Putthasri W. 2014. HIV/AIDS health care challenges for cross-country migrants in low- and middle-income countries: a scoping review. Hiv/Aids 6:19–38.

7. Barnighausen T, Salomon JA, Sangrujee N. 2012. HIV treatment as prevention: issues in economic evaluation. PLoS medicine 9:e1001263.

8. Katlama C, Deeks SG, Autran B, Martinez-Picado J, van Lunzen J, Rouzioux C, Miller M, Vella S, Schmitz JE, Ahlers J, Richman DD, Sekaly RP. 2013. Barriers to a cure for HIV: new ways to target and eradicate HIV-1 reservoirs. Lancet 381:2109–2117.

9. Ho YC, Shan L, Hosmane NN, Wang J, Laskey SB, Rosenbloom DI, Lai J, Blankson JN, Siliciano JD, Siliciano RF. 2013. Replication-competent noninduced proviruses in the latent reservoir increase barrier to HIV-1 cure. Cell 155:540–551.

10. Richman DD, Margolis DM, Delaney M, Greene WC, Hazuda D, Pomerantz RJ. 2009. The challenge of finding a cure for HIV infection. Science 323:1304–1307.

11. Margolis DM. 2014. How Might We Cure HIV? Current infectious disease reports 16:392.

12. Cary DC, Fujinaga K, Peterlin BM. 2016. Molecular mechanisms of HIV latency. The Journal of clinical investigation 126:448–454.

13. Donahue DA, Wainberg MA. 2013. Cellular and molecular mechanisms involved in the establishment of HIV-1 latency. Retrovirology 10:11.

14. Igarashi T, Brown CR, Endo Y, Buckler-White A, Plishka R, Bischofberger N, Hirsch V, Martin MA. 2001. Macrophage are the principal reservoir and sustain high virus loads in rhesus macaques after the depletion of CD4+ T cells by a highly pathogenic simian immunodeficiency virus/HIV type 1 chimera (SHIV): Implications for HIV-1 infections of humans. Proceedings of the National Academy of Sciences of the United States of America 98:658–663.

15. Gartner S, Markovits P, Markovitz DM, Kaplan MH, Gallo RC, Popovic M. 1986. The role of mononuclear phagocytes in HTLV-III/LAV infection. Science 233:215–219.

16. Koenig S, Gendelman HE, Orenstein JM, Dal Canto MC, Pezeshkpour GH, Yungbluth M, Janotta F, Aksamit A, Martin MA, Fauci AS. 1986. Detection of AIDS virus in macrophages in brain tissue from AIDS patients with encephalopathy. Science 233:1089–1093.

17. Honeycutt JB, Wahl A, Baker C, Spagnuolo RA, Foster J, Zakharova O, Wietgrefe S, Caro-Vegas C, Madden V, Sharpe G, Haase AT, Eron JJ, Garcia JV. 2016. Macrophages sustain HIV replication in vivo independently of T cells. The Journal of clinical investigation 126:1353–1366.

18. Lambotte O, Taoufik Y, de Goer MG, Wallon C, Goujard C, Delfraissy JF. 2000. Detection of infectious HIV in circulating monocytes from patients on prolonged highly active antiretroviral therapy. Journal of acquired immune deficiency syndromes 23:114–119.

19. Crowe SM, Sonza S. 2000. HIV-1 can be recovered from a variety of cells including peripheral blood monocytes of patients receiving highly active antiretroviral therapy: a further obstacle to eradication. Journal of leukocyte biology 68:345–350.

20. Price RW, Brew B, Sidtis J, Rosenblum M, Scheck AC, Cleary P. 1988. The brain in AIDS: central nervous system HIV-1 infection and AIDS dementia complex. Science 239:586–592.

21. Gorry PR, Howard JL, Churchill MJ, Anderson JL, Cunningham A, Adrian D, McPhee DA, Purcell DF. 1999. Diminished production of human immunodeficiency virus type 1 in astrocytes results from inefficient translation of gag, env, and nef mRNAs despite efficient expression of Tat and Rev. Journal of virology 73:352–361.

22. Arainga M, Edagwa B, Mosley RL, Poluektova LY, Gorantla S, Gendelman HE. 2017. A mature macrophage is a principal HIV-1 cellular reservoir in humanized mice after treatment with long acting antiretroviral therapy. Retrovirology 14:17.

23. Bisgrove DA, Mahmoudi T, Henklein P, Verdin E. 2007. Conserved P-TEFb-interacting domain of BRD4 inhibits HIV transcription. Proceedings of the National Academy of Sciences of the United States of America 104:13690–13695.

24. Lu P, Qu X, Shen Y, Jiang Z, Wang P, Zeng H, Ji H, Deng J, Yang X, Li X, Lu H, Zhu H. 2016. The BET inhibitor OTX015 reactivates latent HIV-1 through P-TEFb. Scientific reports 6:24100.

25. Pan H, Lu P, Shen Y, Wang Y, Jiang Z, Yang X, Zhong Y, Yang H, Khan IU, Zhou M, Li B, Zhang Z, Xu J, Lu H, Zhu H. 2017. The bromodomain and extraterminal domain inhibitor bromosporine synergistically reactivates latent HIV-1 in latently infected cells. Oncotarget 8:94104–94116.

26. Darcis G, Kula A, Bouchat S, Fujinaga K, Corazza F, Ait-Ammar A, Delacourt N, Melard A, Kabeya K, Vanhulle C, Van Driessche B, Gatot JS, Cherrier T, Pianowski LF, Gama L, Schwartz C, Vila J, Burny A, Clumeck N, Moutschen M, De Wit S, Peterlin BM, Rouzioux C, Rohr O, Van Lint C. 2015. An In-Depth Comparison of Latency-Reversing Agent Combinations in Various In Vitro and Ex Vivo HIV-1 Latency Models Identified Bryostatin-1+JQ1 and Ingenol-B+JQ1 to Potently Reactivate Viral Gene Expression. PLoS pathogens 11:e1005063.

27. Banerjee C, Archin N, Michaels D, Belkina AC, Denis GV, Bradner J, Sebastiani P, Margolis DM, Montano M. 2012. BET bromodomain inhibition as a novel strategy for reactivation of HIV-1. J Leukoc Biol 92:1147–1154.

28. Li G, Nunoya JI, Cheng L, Reszka-Blanco N, Tsao LC, Jeffrey J, Su L. 2017. Regulatory T Cells Contribute to HIV-1 Reservoir Persistence in CD4+ T Cells Through Cyclic Adenosine Monophosphate-Dependent Mechanisms in Humanized Mice In Vivo. J Infect Dis 216:1579–1591.

29. Cheng L, Ma J, Li J, Li D, Li G, Li F, Zhang Q, Yu H, Yasui F, Ye C, Tsao LC, Hu Z, Su L, Zhang L. 2017. Blocking type I interferon signaling enhances T cell recovery and reduces HIV-1 reservoirs. The Journal of clinical investigation 127:269–279.

30. Sattentau QJ, Stevenson M. 2016. Macrophages and HIV-1: An Unhealthy Constellation. Cell host & microbe 19:304–310.

31. Calvanese V, Chavez L, Laurent T, Ding S, Verdin E. 2013. Dual-color HIV reporters trace a population of latently infected cells and enable their purification. Virology 446:283–292.

32. Chavez L, Calvanese V, Verdin E. 2015. HIV Latency Is Established Directly and Early in Both Resting and Activated Primary CD4 T Cells. PLoS pathogens 11:e1004955.

33. Wei DG, Chiang V, Fyne E, Balakrishnan M, Barnes T, Graupe M, Hesselgesser J, Irrinki A, Murry JP, Stepan G, Stray KM, Tsai A, Yu H, Spindler J, Kearney M, Spina CA, McMahon D, Lalezari J, Sloan D, Mellors J, Geleziunas R, Cihlar T. 2014. Histone deacetylase inhibitor romidepsin induces HIV expression in CD4 T cells from patients on suppressive antiretroviral therapy at concentrations achieved by clinical dosing. PLoS pathogens 10:e1004071.

34. Rasmussen TA, Schmeltz Sogaard O, Brinkmann C, Wightman F, Lewin SR, Melchjorsen J, Dinarello C, Ostergaard L, Tolstrup M. 2013. Comparison of HDAC inhibitors in clinical development: effect on HIV production in latently infected cells and T-cell activation. Human vaccines & immunotherapeutics 9:993–1001.

35. Archin NM, Liberty AL, Kashuba AD, Choudhary SK, Kuruc JD, Crooks AM, Parker DC, Anderson EM, Kearney MF, Strain MC, Richman DD, Hudgens MG, Bosch RJ, Coffin JM, Eron JJ, Hazuda DJ, Margolis DM. 2012. Administration of vorinostat disrupts HIV-1 latency in patients on antiretroviral therapy. Nature 487:482–485.

36. Ammosova T, Berro R, Kashanchi F, Nekhai S. 2005. RNA interference directed to CDK2 inhibits HIV-1 transcription. Virology 341:171–178.

37. Debebe Z, Ammosova T, Breuer D, Lovejoy DB, Kalinowski DS, Kumar K, Jerebtsova M, Ray P, Kashanchi F, Gordeuk VR, Richardson DR, Nekhai S. 2011. Iron chelators of the di-2-pyridylketone thiosemicarbazone and 2-benzoylpyridine thiosemicarbazone series inhibit HIV-1 transcription: identification of novel cellular targets--iron, cyclin-dependent kinase (CDK) 2, and CDK9. Molecular pharmacology 79:185–196.

38. Agbottah E, de La Fuente C, Nekhai S, Barnett A, Gianella-Borradori A, Pumfery A, Kashanchi F. 2005. Antiviral activity of CYC202 in HIV-1-infected cells. The Journal of biological chemistry 280:3029–3042.

39. Salerno D, Hasham MG, Marshall R, Garriga J, Tsygankov AY, Grana X. 2007. Direct inhibition of CDK9 blocks HIV-1 replication without preventing T-cell activation in primary human peripheral blood lymphocytes. Gene 405:65–78.

40. Medina-Moreno S, Dowling TC, Zapata JC, Le NM, Sausville E, Bryant J, Redfield RR, Heredia A. 2017. Targeting of CDK9 with indirubin 3’-monoxime safely and durably reduces HIV viremia in chronically infected humanized mice. PLoS One 12:e0183425.

41. Okamoto M, Hidaka A, Toyama M, Hosoya T, Yamamoto M, Hagiwara M, Baba M. 2015. Selective inhibition of HIV-1 replication by the CDK9 inhibitor FIT-039. Antiviral research 123:1–4.

42. Jerebtsova M, Kumari N, Xu M, de Melo GB, Niu X, Jeang KT, Nekhai S. 2012. HIV-1 Resistant CDK2-Knockdown Macrophage-Like Cells Generated from 293T Cell-Derived Human Induced Pluripotent Stem Cells. Biology 1:175–195.

43. Alexander LT, Mobitz H, Drueckes P, Savitsky P, Fedorov O, Elkins JM, Deane CM, Cowan-Jacob SW, Knapp S. 2015. Type II Inhibitors Targeting CDK2. ACS chemical biology 10:2116–2125.

44. Thompson KA, Cherry CL, Bell JE, McLean CA. 2011. Brain cell reservoirs of latent virus in presymptomatic HIV-infected individuals. The American journal of pathology 179:1623–1629.

45. Hellmuth J, Valcour V, Spudich S. 2015. CNS reservoirs for HIV: implications for eradication. Journal of virus eradication 1:67–71.

46. Brenchley JM, Hill BJ, Ambrozak DR, Price DA, Guenaga FJ, Casazza JP, Kuruppu J, Yazdani J, Migueles SA, Connors M, Roederer M, Douek DC, Koup RA. 2004. T-cell subsets that harbor human immunodeficiency virus (HIV) in vivo: implications for HIV pathogenesis. Journal of virology 78:1160–1168.

47. Buzon MJ, Sun H, Li C, Shaw A, Seiss K, Ouyang Z, Martin-Gayo E, Leng J, Henrich TJ, Li JZ, Pereyra F, Zurakowski R, Walker BD, Rosenberg ES, Yu XG, Lichterfeld M. 2014. HIV-1 persistence in CD4+ T cells with stem cell-like properties. Nature medicine 20:139–142.

48. Chomont N, El-Far M, Ancuta P, Trautmann L, Procopio FA, Yassine-Diab B, Boucher G, Boulassel MR, Ghattas G, Brenchley JM, Schacker TW, Hill BJ, Douek DC, Routy JP, Haddad EK, Sekaly RP. 2009. HIV reservoir size and persistence are driven by T cell survival and homeostatic proliferation. Nature medicine 15:893–900.

49. Jorajuria S, Dereuddre-Bosquet N, Becher F, Martin S, Porcheray F, Garrigues A, Mabondzo A, Benech H, Grassi J, Orlowski S, Dormont D, Clayette P. 2004. ATP binding cassette multidrug transporters limit the anti-HIV activity of zidovudine and indinavir in infected human macrophages. Antiviral therapy 9:519–528.

50. Zha W, Wang G, Xu W, Liu X, Wang Y, Zha BS, Shi J, Zhao Q, Gerk PM, Studer E, Hylemon PB, Pandak WM, Jr., Zhou H. 2013. Inhibition of P-glycoprotein by HIV protease inhibitors increases intracellular accumulation of berberine in murine and human macrophages. PloS one 8:e54349.

51. Choo EF, Leake B, Wandel C, Imamura H, Wood AJ, Wilkinson GR, Kim RB. 2000. Pharmacological inhibition of P-glycoprotein transport enhances the distribution of HIV-1 protease inhibitors into brain and testes. Drug metabolism and disposition: the biological fate of chemicals 28:655–660.

52. Lorenzo-Redondo R, Fryer HR, Bedford T, Kim EY, Archer J, Kosakovsky Pond SL, Chung YS, Penugonda S, Chipman JG, Fletcher CV, Schacker TW, Malim MH, Rambaut A, Haase AT, McLean AR, Wolinsky SM. 2016. Persistent HIV-1 replication maintains the tissue reservoir during therapy. Nature 530:51–56.

53. Gendelman HE, Orenstein JM, Martin MA, Ferrua C, Mitra R, Phipps T, Wahl LA, Lane HC, Fauci AS, Burke DS, et al. 1988. Efficient isolation and propagation of human immunodeficiency virus on recombinant colony-stimulating factor 1-treated monocytes. The Journal of experimental medicine 167:1428–1441.

54. Carter CA, Ehrlich LS. 2008. Cell biology of HIV-1 infection of macrophages. Annual review of microbiology 62:425–443.

55. Wei P, Garber ME, Fang SM, Fischer WH, Jones KA. 1998. A novel CDK9-associated C-type cyclin interacts directly with HIV-1 Tat and mediates its high-affinity, loop-specific binding to TAR RNA. Cell 92:451–462.

56. Breuer D, Kotelkin A, Ammosova T, Kumari N, Ivanov A, Ilatovskiy AV, Beullens M, Roane PR, Bollen M, Petukhov MG, Kashanchi F, Nekhai S. 2012. CDK2 regulates HIV-1 transcription by phosphorylation of CDK9 on serine 90. Retrovirology 9:94.

57. Ammosova T, Berro R, Jerebtsova M, Jackson A, Charles S, Klase Z, Southerland W, Gordeuk VR, Kashanchi F, Nekhai S. 2006. Phosphorylation of HIV-1 Tat by CDK2 in HIV-1 transcription. Retrovirology 3:78.

58. Pauls E, Ruiz A, Badia R, Permanyer M, Gubern A, Riveira-Munoz E, Torres-Torronteras J, Alvarez M, Mothe B, Brander C, Crespo M, Menendez-Arias L, Clotet B, Keppler OT, Marti R, Posas F, Ballana E, Este JA. 2014. Cell cycle control and HIV-1 susceptibility are linked by CDK6-dependent CDK2 phosphorylation of SAMHD1 in myeloid and lymphoid cells. J Immunol 193:1988–1997.

59. Jiang Q, Zhang L, Wang R, Jeffrey J, Washburn ML, Brouwer D, Barbour S, Kovalev GI, Unutmaz D, Su L. 2008. FoxP3+CD4+ regulatory T cells play an important role in acute HIV-1 infection in humanized Rag2−/−gammaC−/− mice in vivo. Blood 112:2858–2868.

60. Li G, Cheng M, Nunoya J, Cheng L, Guo H, Yu H, Liu YJ, Su L, Zhang L. 2014. Plasmacytoid dendritic cells suppress HIV-1 replication but contribute to HIV-1 induced immunopathogenesis in humanized mice. PLoS pathogens 10:e1004291.

61. Oswald-Richter K, Grill SM, Shariat N, Leelawong M, Sundrud MS, Haas DW, Unutmaz D. 2004. HIV infection of naturally occurring and genetically reprogrammed human regulatory T-cells. PLoS biology 2:E198.

62. Zhang L, Jiang Q, Li G, Jeffrey J, Kovalev GI, Su L. 2011. Efficient infection, activation, and impairment of pDCs in the BM and peripheral lymphoid organs during early HIV-1 infection in humanized rag2(-)/(-)gamma C(-)/(-) mice in vivo. Blood 117:6184–6192.

63. Zhang L, Kovalev GI, Su L. 2007. HIV-1 infection and pathogenesis in a novel humanized mouse model. Blood 109:2978–2981.

64. Halper-Stromberg A, Lu CL, Klein F, Horwitz JA, Bournazos S, Nogueira L, Eisenreich TR, Liu C, Gazumyan A, Schaefer U, Furze RC, Seaman MS, Prinjha R, Tarakhovsky A, Ravetch JV, Nussenzweig MC. 2014. Broadly neutralizing antibodies and viral inducers decrease rebound from HIV-1 latent reservoirs in humanized mice. Cell 158:989–999.

65. Livak KJ, Schmittgen TD. 2001. Analysis of relative gene expression data using real-time quantitative PCR and the 2(-Delta Delta C(T)) Method. Methods 25:402–408.

